# Mutation of *M. truncatula SOBIR1* affects rhizobial specificity and arbuscular mycorrhizal colonization

**DOI:** 10.1101/2025.09.17.676642

**Authors:** E. Schnabel, L. Müller, C.I. Pislariu, S. Ivanov, H. Samara, M.J. Harrison, J. Frugoli

## Abstract

Legume plants form symbiotic interactions with bacteria (rhizobia sp.) to obtain fixed nitrogen and arbuscular mycorrhizal fungi to obtain phosphorus and other nutrients. These interactions require the plant to distinguish beneficial from pathogenic organisms and trigger the plant’s innate immune system. We identified mutants in the *Medicago truncatula SOBIR1* gene, known to be involved in innate immune response in multiple plants. We examined nodulation with multiple strains and species of *Sinorhizobium* and examined mycorrhizal interactions with the arbuscular mycorrhizal fungus *Glomus versiforme (recently renamed Diversispora epigaea)*. Plants containing mutations in *SOBIR1* exhibit normal nodulation with strains of *S. meliloti*, but fewer nodules when inoculated with *S. medicae* strains. The *S. medicae* nodules show evidence of accumulation of polyphenolic compounds, and abnormal arrangement of the rhizobia within the nodules. In contrast, when inoculated with *Glomus versiforme. M. truncatula sobir1* mutant plants have increased mycorrhizal colonization and arbuscule number compared to the wild type. We localized a tagged SOBIR1 protein to the periarbuscular membrane interface. Together the data suggest the immune kinase SOBIR1 is involved in both nodulation and mycorrhizal interactions, but the effects of mutation differ.

## Introduction

Nitrogen is crucial to plant growth, and while it is the most abundant gas in the Earth’s atmosphere, it can only be used by plants when the triple bond of the gas is broken. Most organisms on the planet do not have the ability to break the bond, but some prokaryotic organisms (diazotrophs) contain an enzyme, nitrogenase, that can break the bond and produce forms of nitrogen usable to plants: NO_3_^-^ and NH_4_^+^. This biological nitrogen fixation is the largest contributor of atmospheric nitrogen to the biosphere (Herridge et al., 2008).

Legume plants form a symbiotic association with rhizobia bacteria, one of the nitrogen-fixing diazotrophs, creating specialized plant organs called nodules to house the bacteria within the roots of the plant. This allows legumes to grow in nitrogen-poor soil by using the nitrogen fixed by the bacteria in exchange for carbon skeletons from photosynthesis. The interspecies symbiosis is species limited with complex regulation controlled by many genes in both organisms. In plants, these genes regulate whether the rhizobia are compatible, whether the plant needs nitrogen and thus can benefit from the symbiosis, and how many nodules to form if the plant allows the symbiosis (Roy et al., 2020; Chaulagain and Frugoli, 2021).

Since plants interact with multiple microbes at the interface between the root and the soil, distinguishing between beneficial bacteria and fungi and pathogenic bacteria and fungi, as well as benign endophytes is critical to successful symbioses. Most plants have an innate immune system to recognize and respond to these bacteria and fungi (Ngou et al., 2022). Indeed, legumes responding to beneficial rhizobia initiate a brief defense response, which the rhizobia suppress in order to infect and form nodules (reviewed in (Grundy et al., 2023)).

Although more broadly distributed to roughly 80% of land plants, legumes also engage in symbiosis with arbuscular mycorrhizal fungi (AM symbiosis) which involves mycorrhizal fungi forming structures within plant root cells and delivering nutrients through the external hyphal network of the fungi, such as phosphorus and nitrogen in exchange for carbon (Smith and Read, 2008). While not as species-limited as the rhizobial symbiosis, the plants still distinguish between pathogenic and beneficial fungi and mount a defense response, which is then suppressed by symbiotic fungi, including a switch that can modulate the response based on environmental conditions (Zhang et al., 2021). Many of the same molecular steps in immune response involved in the rhizobial symbiosis also occur in the response to AM fungi (reviewed in (Díaz et al., 2025)).

Since both rhizobial symbiosis and AM symbiosis involve suppression of the immune response to rhizobial or mycorrhizal partners, we reasoned that some receptor kinases identified as affecting immunity and development might be candidates for controlling the early establishment of these interactions. We wondered if the *SOBIR1* gene, identified as part of the immune response in Arabidopsis, might be involved in these two symbioses in legumes. Arabidopsis SOBIR1 constitutively associates with multiple Leucine-Rich Repeat Receptor-Like Proteins (LRR-RLPs), providing the essential intracellular kinase domain that RLPs lack (Gust and Felix, 2014) and is involved in a molecular complex initiating pattern-triggered immunity in response to microbial necrosis- and ethylene-inducing proteins (Albert et al., 2015). SOBIR1 is also involved in the response to pathogenic fungi. In *Solanum* species, SOBIR1 and SOBIR1-like are required for resistance against *Phytophthora infestans* and *Hyaloperonospora arabidopsidis* (Domazakis et al., 2018) and in tomato, the Fusarium wilt resistance gene depends on SOBIR1 to signal (Catanzariti et al., 2017).

To investigate the role of *MtSOBIR1*, we isolated two independent mutant alleles in the *M. truncatula SOBIR1* gene (*Medtr3g075440*) and tested nodule number phenotype using multiple strains of rhizobia, adding to similar recent work which used *M. truncatula* mutants but only two strains of rhizobia (Sarrette et al., 2025). We also tested mycorrhizal interactions with *Glomus versiforme*. Our results show that even accounting for slightly different rhizobial responses in different ecotypes of *M. truncatula, sobir1* mutants appear to have a differential response to *S. medicae* versus *S. meliloti*, resulting in few nodules when inoculated with *S. medicae* strains, evidence of accumulation of polyphenolic compounds, and abnormal arrangement of the rhizobia within the nodules. In contrast, investigations into mycorrhizal interactions revealed *sobir1* mutant plants have increased mycorrhizal colonization suggesting a role in this symbiosis as well, but with opposite effects from nodulation. We show that the SOBIR1 protein localizes to the periarbuscular and plasma membrane during the mycorrhizal interaction and together these observations suggest different roles of SOBIR1 in immune response and in nodule formation and structure, versus arbuscular number and AM fungal root colonization.

## Results and Discussion

After isolating two independent homozygous mutants in the *SOBIR1* gene of *M. truncatula* (see Methods), we identified the location of the *Tnt1* insertions in each mutant (Figure 1A). Both mutants contain insertions at the beginning of the gene; the insertion in *sobir1-1* is within nucleotides encoding the signal peptide, while the insertion in *sobir1-2* is located in the ectodomain domain upstream of the leucine-rich repeats (Figure 1A).

**Figure 1.**
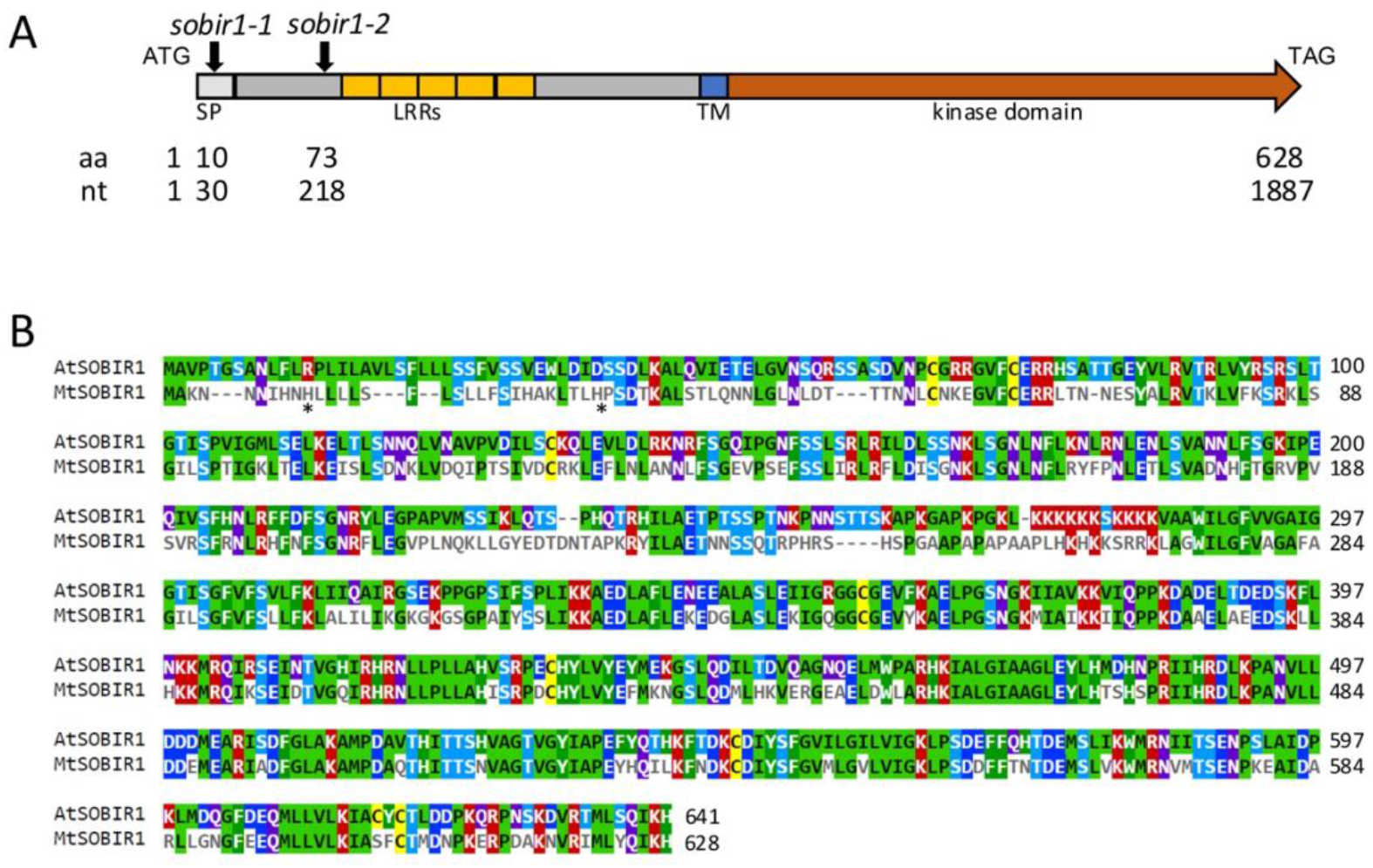
Molecular identification of two mutant alleles of *MtSOBIR1*, and comparison to the Arabidopsis protein. (A) Structure of MtSOBIR1 and location of insertions in mutant alleles. MtSOBIR1, encoded by *Medtr3g075440*, is a predicted 628 amino acid receptor-like kinase with a cleaved signal peptide (SP), extracellular domain with leucine-rich repeats (LRRs), transmembrane domain (TM), and cytoplasmic kinase domain. Two mutant alleles, *sobir1-1* and *sobir1-2*, have Tnt1 insertions near the beginning of the coding sequence at nucleotide positions 30 and 218, respectively. (B) Alignment of MtSOBIR1 and AtSOBIR1 with Clustal Omega visualized with mView. Asterisks indicate location of insertions. Residues in MtSOBIR1 that are identical to AtSOBIR1 are colored on both lines. The colors refer to amino acid properties. Red indicates small, hydrophobic, or aromatic residues, blue=acidic residues, magenta=basic residues and green=hydroxyl, sulfhydryl, or amine-containing residues.

Given their locations, both insertions likely disrupt localization to the membrane or ligand binding, but the kinase domain may be active. This is important because the kinase domain is required for immune response in Arabidopsis (Van Der Burgh et al., 2019). Comparing the *M. truncatula* gene to Arabidopsis *SOBIR1* using mView (Madeira et al., 2019), *MtSOBIR1* is 13 amino acids shorter than the Arabidopsis gene. Most of the “missing” amino acids are early in the gene near the insertions. As might be expected for functional domains, the transmembrane domain and kinase domains are highly similar between the two species.

We tested the nodule number phenotype of the mutants using multiple strains of rhizobia. Most of the forward genetic mutants in *M. truncatula* were created in the A17 ecotype used for the public genome sequence. In our aeroponic system described in Methods, A17 plants nodulate well with the ABS7 strain of *S. medicae*, producing 7-10 nodules per plant at 12 days post inoculation (dpi). However, the A17 ecotype is recalcitrant to transformation due to a genome rearrangement (Kamphuis et al., 2007); therefore the *Tnt1* insertion lines were created in the R108 ecotype. R108 is relatively easy to transform but is genetically different enough from most other *M. truncatula* ecotypes that it has been proposed as a different species, *M*.*littoralis* (Choi et al., 2022). While R108 nodulates when inoculated with *S. medicae* ABS7, the average number of nodules per plant at 12 dpi is 3-7 in our aeroponic system. Many labs working with *M. truncatula* use strains of *S. meliloti* for inoculation of both the A17 and R108 ecotypes of *M. truncatula*, as this strain has a long history of molecular genetics, the advantage of multiple rhizobial mutants, and produces similar nodulation in the two ecotypes.

Therefore, we chose to inoculate wild-type R108 plants and the *sobir1-1* and *sobir1-2* mutants in the R108 background with two strains of *S. medicae* and three strains of *S. meliloti* in an aeroponic chamber and counted nodules at 12 dpi. Compared to wild type (R108), both *sobir1* alleles displayed significantly reduced nodule number with the two *S. medicae* strains (Figure 2A). When inoculated with any of the three strains of *S. meliloti*, the *sobir1* mutants were not significantly different from the wild type inoculated with the same strain (Figure 2B). This suggests that the reduced nodulation in the *SOBIR1*mutants is not strain-specific but depends on the species of rhizobia.

**Figure 2.**
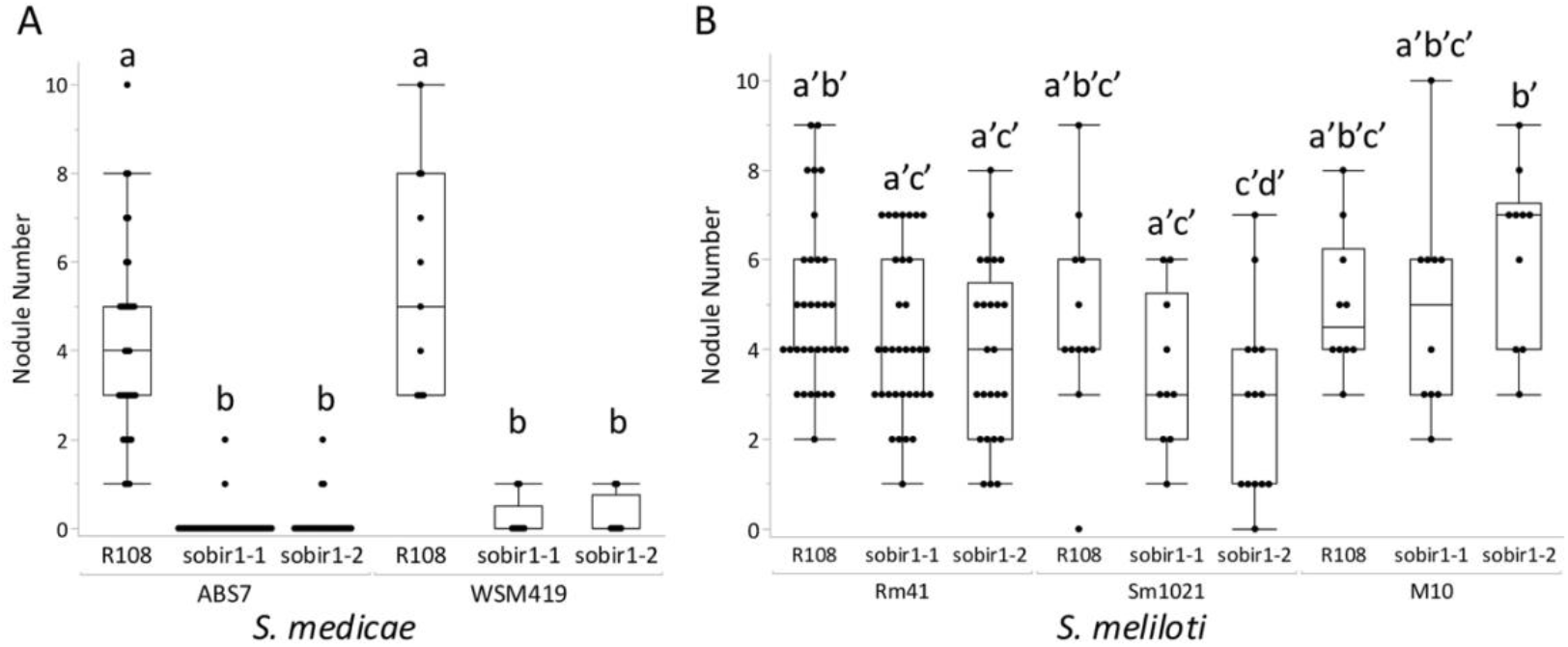
Disruption of *SOBIR1* impacts nodulation by rhizobia in a strain-specific manner. Nodulation of *M. truncatula* R108 (wild type) and two *sobir1* alleles was assessed 12 days after inoculation with (A) *Sinorhizobium medicae* strains ABS7 and WSM419 and (B) *S. meliloti* strains Rm41, Sm1021, and M10. Nodulation in both *sobir1* mutants was significantly lower than in wild type with both *S. medicae* strains, while no difference was observed with *S. meliloti* strains (Kruskal-Wallis test with Bonferroni correction, p < 0.05, samples with no significant difference are indicated with a similar letter).

We also counted nodules at 21 days post-inoculation, and the numbers did not change (data not shown) demonstrating that *sobir1* mutants are not slow to grow nodules with *S. medicae*; rather, the mutants rarely form nodules. Examination of the few nodules that did form revealed visual defects in the nodules (Figure 3). At 12 dpi, the point at which we counted nodules in Figure 1, the nodules on *sobir1* mutants inoculated with *S. medicae* ABS7 are not round, but flattened, barely emerged from the root, and not visibly pink (Figure 3A). The pink color observed in wild type comes from leghemoglobin, a molecule necessary for nitrogen fixation. Observed at 20 dpi, the indeterminant nodules on the R108 wild type are a healthy pink and elongating, while the nodules on the *sobir1* mutants have not yet grown. They have also not turned pink and seem to be accumulating polyphenols (indicated by the brown color), suggesting necrosis or stress, a phenotype associated with incompatible rhizobia interactions (Vasse et al., 1993). However, when inoculated with *S. meliloti* Rm41 (Figure 3B), nodulation in both mutants is indistinguishable from wild type at 20 dpi in our observations.

**Figure 3.**
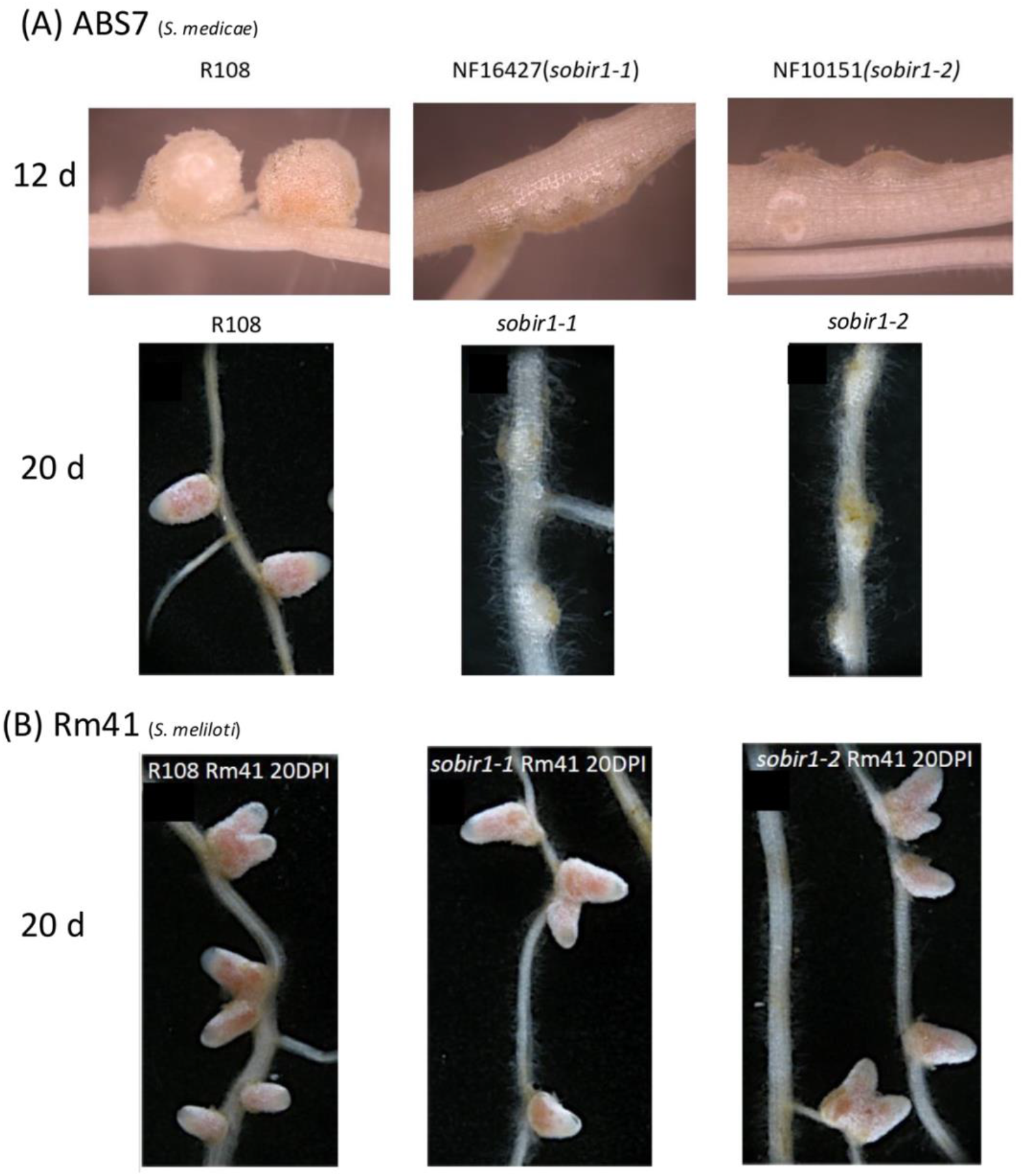
Macroscopic phenotype depends on species of rhizobia. (A) Macroscopic phenotype of *sobir1* mutants compared to R108 wild type when inoculated *with S. medicae* (ABS7) (B) Macroscopic phenotype of *sobir1* mutants compared to R108 wild type when inoculated with *S. meliloti* (Rm41).

Rhizobial strains Rm41 and ABS7 used carry a LacZ marker, allowing examination of the arrangement of rhizobia within the nodules by staining. In a mature indeterminant nodule formed by *M. truncatula*, the nodule is divided into zones with different morphologies. Zone 1 is the meristematic zone at the tip of the nodules, which continues to elongate. Behind it is Zone II, the zone of infection and differentiation. Here, rhizobia are dividing and infecting the plant cells. In Zone III, the differentiated rhizobia fix nitrogen, and in Zone IV, next to the root, the senescence zone in which bacteria die after several weeks and are degraded (Vasse et al., 1990). At 20 dpi, R108 and *Mtsobir1-2* nodules display the normal pattern of zones, with a minimal senescence zone (Figure 4 A, C). However, when inoculated with ABS7, *Mtsobir1-2* mutants display a disorganized nodule structure (Figure 4B). The nodule shape is more round than cylindrical, and zonation is disrupted; a well-defined proximal zone II characterized by high symbiosome proliferation, seen as a dark blue band in Fig. 4 A and C, is missing in *Mtsobir1-2* nodules harboring ABS7. Even in the proximal zone (at the bottom of Figure 4B), infected cells determined by the LacZ stain have multiple vacuoles, so that, instead of seeing a “ring” of bacteroids surrounding a central vacuole, the intense blue color has a patchy distribution within cells. The misshapen cells could also be due to generalized damage resulting from altered membrane integrity. Vacuolar fragmentation or convolution has been shown to occur during stress (reviewed by (Zhang et al., 2014), reinforcing the possibility that this is an immune response. The observation of disorganization was reinforced by confocal microscopy of individual cells within the nodules (Figure 5). In R108 plants inoculated with Rm41 rhizobia (Figure 5A) or ABS7 rhizobia (Figure 5B), the rhizobial bacteroids within a cell (stained green) surround a central vacuole. In contrast, while *sobir1-2* plants inoculated with Rm41 show the same organization (Figure 5C), when inoculated with ABS7 rhizobia the bacteroids inside nodules in *Mtsobir1-2* plants are scattered throughout the cell and seem to occasionally form multiple foci. They are also less defined and minimally elongated, in contrast to Rm41 in *Mtsobir1-2* nodules. This random distribution of bacteroids inside infected cells may be due to multiple smaller vacuoles. In compatible interactions, host cell nuclei are rather inconspicuous, appearing as a pale fluorescent structure with smooth edges and lacking the brightness of the rhizobia (Figure 5B). In *Mtsobir1-2* nodules infected with ABS7, the nuclei are enlarged and misshapen with rough edges and condensed chromatin visible as brightly fluorescent speckles (Figure 5 D). These cellular modifications are indicative of a stress response to ABS7 rhizobia.

**Figure 4.**
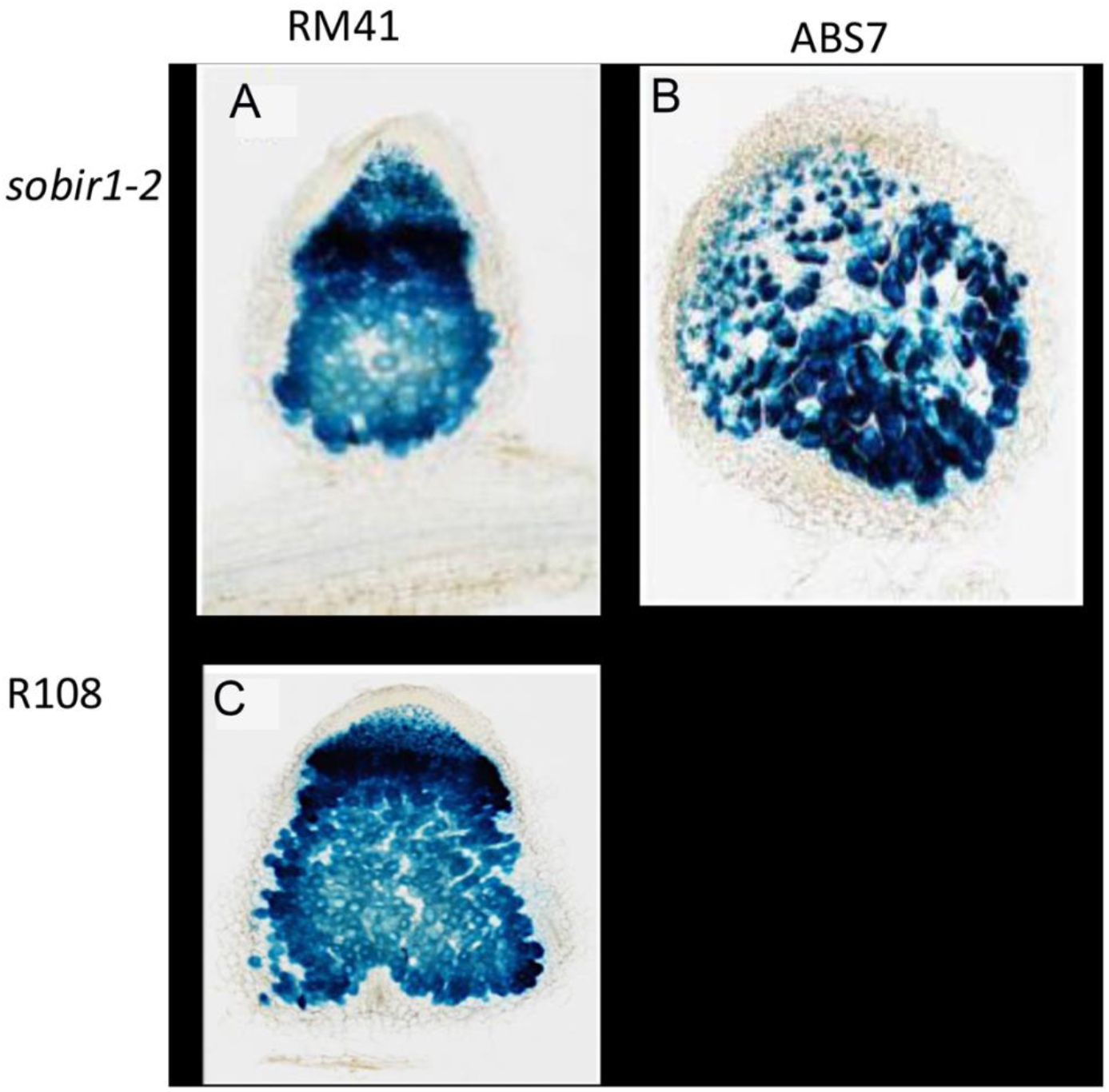
LacZ staining of 20 dpi nodules in *sobir1* mutants compared to R108 wild type when inoculated *with S. medicae* (ABS7) versus *S. meliloti* (Rm41). (A) LacZ staining of *Mtsobir 1-2* nodule inoculated with Rm41. (B) LacZ staining of R108 nodule inoculated with Rm41. (C) LacZ staining of *Mtsobir1-2* nodule inoculated with ABS7.

**Figure 5.**
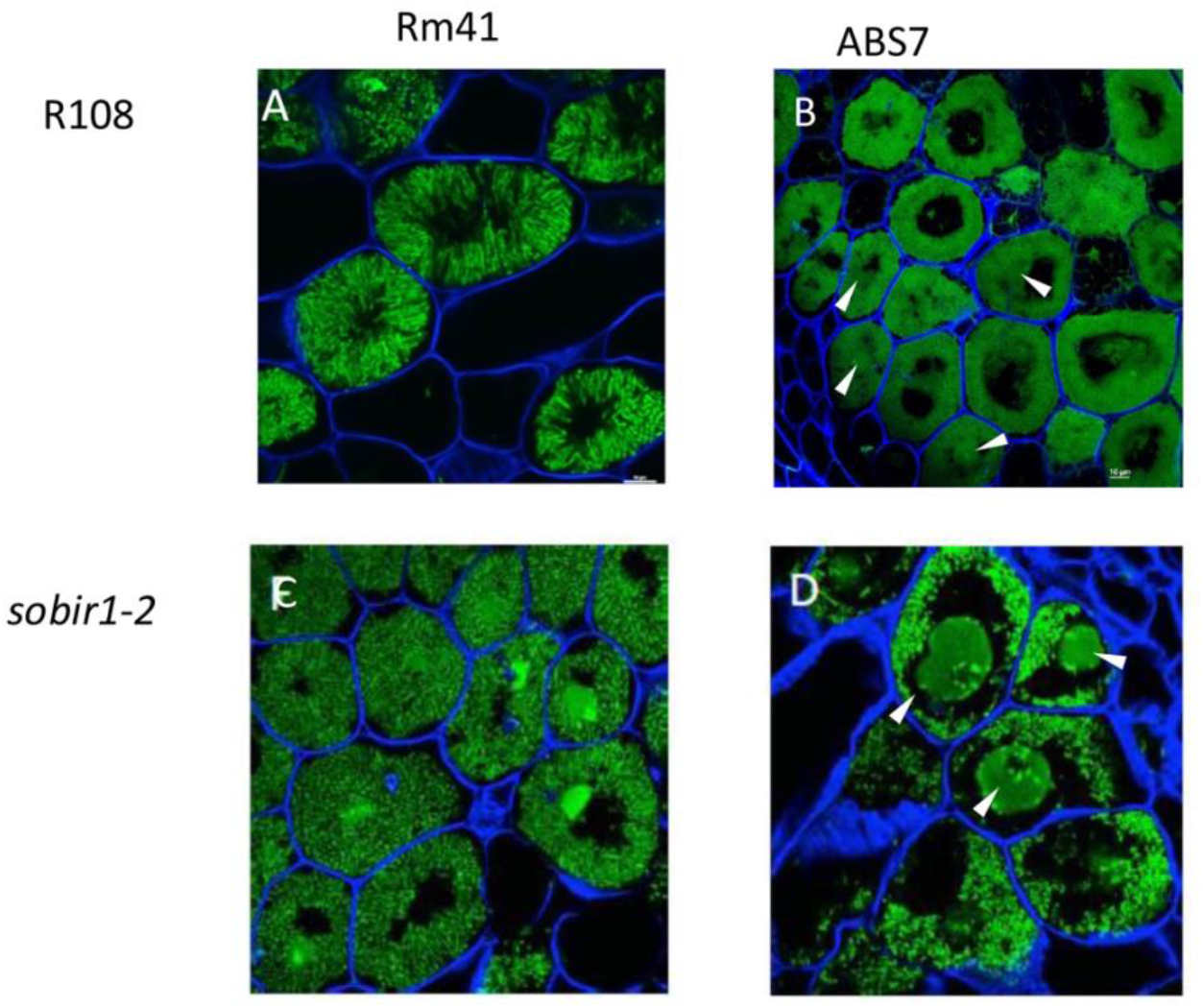
Confocal microscopy of 12 dpi nodules in *sobir1* mutants compared to R108 wild type when inoculated *with S. medicae* (ABS7) versus *S. meliloti* (Rm41). Plants were stained with Syto13 for DNA (green fluorescence) and calcofluor white (blue fluorescence) for plant cell walls. White arrows indicate cell nuclei. (A) Cells from nodules on R108 plants inoculated with RM41 or (B) ABS7. (C) Cells from nodules on *Mtsobir 1-2* plants inoculated with RM41 or (D) ABS7.

We reasoned that if nodule development was affected in *M. truncatula sobir1* mutants, arbuscular mycorrhizal (AM) symbiosis might also be affected, since both interactions share a set of genetic components (the common symbiotic pathway (Genre and Russo, 2016)) and both involve microbes; the plant immune system must gatekeep by distinguishing between pathogenic and beneficial interactions. Interestingly, in contrast to the reduced nodule number observed with some strains of rhizobia, M. *truncatula sobir1* mutants displayed a higher percentage of root colonization (Figure 6A) and higher numbers of arbuscules relative to R108 wild-type roots when colonized with the AM fungus *Glomus versiforme* (Figure 6B, C). This result suggests the immune response triggered by the initiation of rhizobial symbiosis and AM symbiosis, at least with *G. versiforme*, occurs upstream or downstream of the Common Symbiotic Pathway, with different effects (decreased infection for rhizobia, increased for *G. versiforme*).

**Figure 6.**
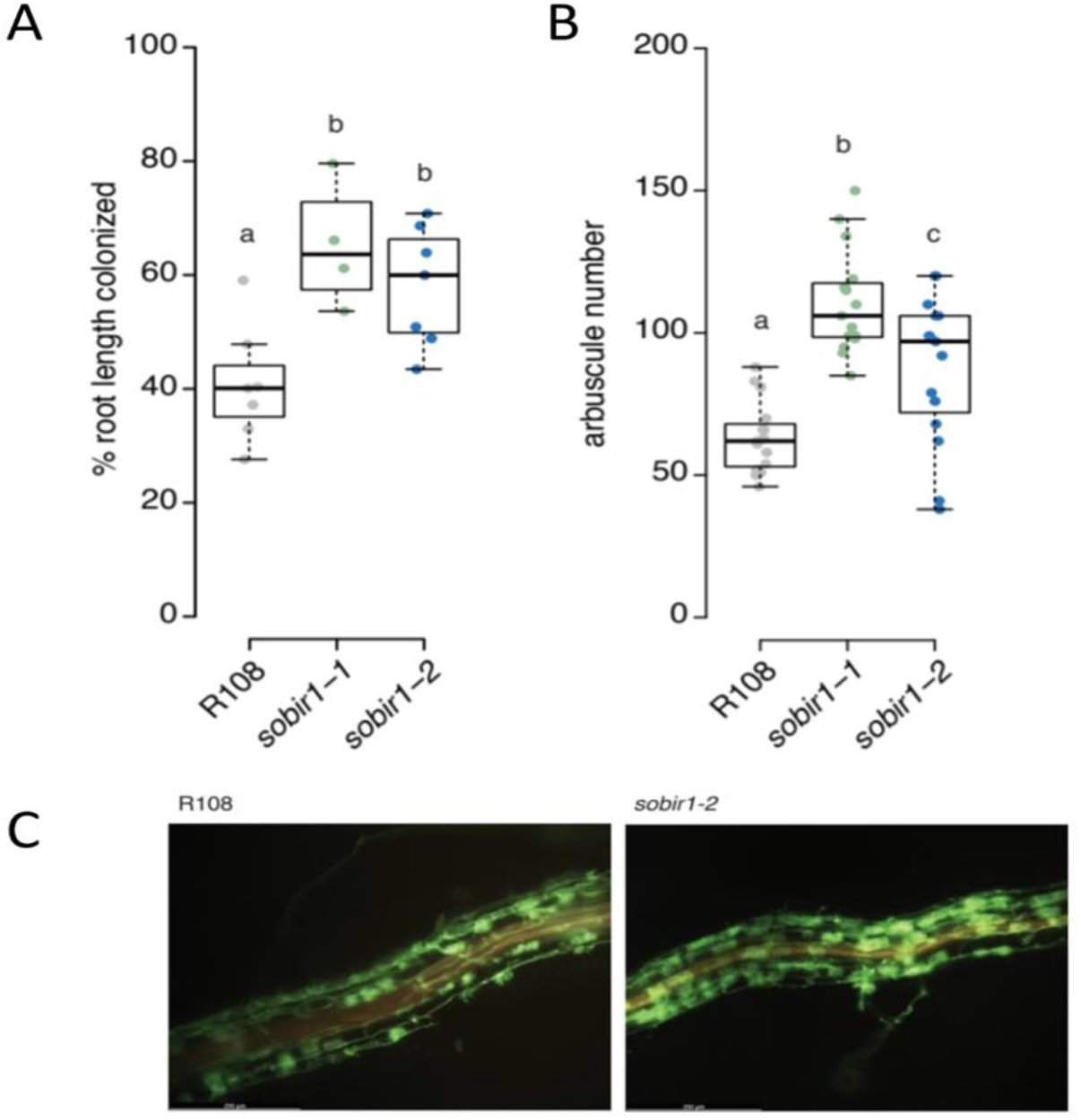
*Mtsobir1* mutants are hyperinfected by *Glomus versiforme*. Colonization of *M. truncatula* roots with the AM fungus *G. versiforme* was assessed 4 weeks after infection as (A) extent of root length colonization and (B) number of arbuscules formed in R108 (wild type) and two *Mtsobir-1* mutants. Data is represented as box-and-whiskers plots showing upper and lower quartiles, and the median as a horizontal black line. Colored points represent individual measurements. Both mutants showed significantly higher interaction than R108 by a Kruskal-Wallis test followed by a two-sided Dunn’s post-hoc test with multiple comparison correction after the Benjamini-Hochberg method. Pairwise differences were considered significant if p<0.05, denoted by different letters in the graphs. All statistical analyses were performed using R software. (C) Excess colonization of *sobir1-2* versus R108 visualized by WGA-Alexafluor488. (see Methods).

To further explore the SOBIR1 protein in AM interactions, we localized the SOBIR1 protein in wild-type plants undergoing mycorrhizal symbiosis. Using a cpVenus-tagged protein in plants that also express a membrane marker from an AM-inducible promoter, we observed co-localization of tagged SOBIR1 with both the periarbuscular membrane and plasma membrane (Figure 7). This suggests the effect of mutation of SOBIR1 on mycorrhizal interactions (and perhaps nodulation) may involve disruption of signaling at a membrane interface.

**Figure 7.**
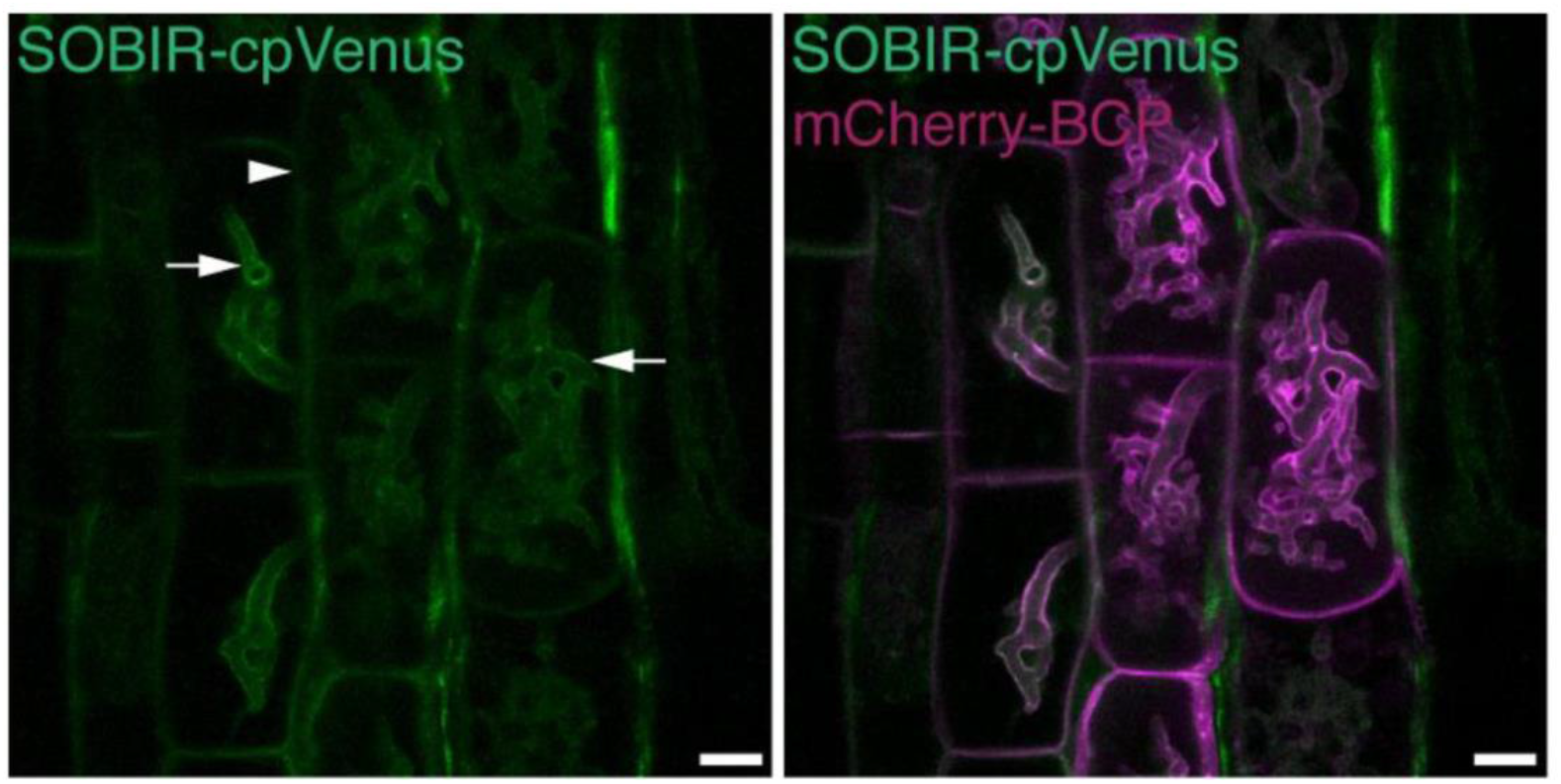
Sub-cellular location of SOBIR1-cpVenus in *M. truncatula* root cortical cells containing arbuscules of AM fungus *Rhizophagus irregularis*. SOBIR1-cpVenus was visible on the periarbuscular membrane (arrow) and plasma membrane (arrowhead). The periarbuscular membrane and plasma membrane are marked by mCherry-BCP. Confocal microscopy images (z-stack projection, n=10 of optical slices, 0.5 µm interval). *n* – nucleus. Scale bars, 10µm.

## Conclusions

Our data show that a functional copy of SOBIR1 is necessary for normal nodule formation in R108 *M. truncatula* plants inoculated with strains of *S. medicae*, the natural symbiont of *M. truncatula*, but not *S. meliloti*, the natural symbiont of *M. sativa*. In contrast, mutation of *SOBIR1* in *M. truncatula* increases colonization of the AM fungus *Glomus versiforme*, and the protein appears to localize to the plant/fungal interface.

## Methods

### Plant growth conditions for rhizobia experiments Figure 2 and Figure 3A

Seeds were germinated as described previously (Schnabel et al., 2010). This involves scarification in concentrated sulfuric acid, imbibition in water, vernalization at 4°C for 2 days, followed by overnight germination at room temperature in the dark. One-day-old seedlings were placed in an aeroponic chamber and grown at 21°C-25°C; 14h/10h light/dark cycle and inoculated as described in (Cai et al., 2023). Depending on the experiment plants were inoculated with *S. meliloti* RM41 (NCBI:txid1230587), 1021 (NCBI:txid266834), M10 (NCBI:txid1230602), *S. medicae* ABS7 (Leong et al. 1985) or WSM419 (NCBI:txid366394) four days post germination. Nodule count was performed 12 days post-inoculation (dpi), unless otherwise noted. For seed collection and genetic crosses, plants were grown in a greenhouse with supplemental light on a 14h/10h light/dark cycle at 21°C-25°C

### Identification and Isolation of Mutants

Two pools of *Tnt1* insertion mutants carrying an insertion in *MtSOBIR1* were digitally identified in the *M. truncatula Tnt1* insertion library database (Tadege et al., 2008; Pislariu et al., 2012) as carrying an insertion in *MtSOBIR1* were screened by PCR for insertions in the gene. From each pool 4-6 plants were grown, and PCR was used to identify individual plants either homozygous or heterozygous for an insertion utilizing gene-specific primers (JF3067 or JF3068; Table 1) near the putative insertion sites paired with a Tnt1-specific primer (Tntail3). The NF16427 pool contained a plant homozygous for an insertion (*sobir1-1*), and the NF10151 pool contained a plant heterozygous for an insertion in a different location (*sobir1-2*). The exact locations of the insertions were determined by sequencing the PCR products. Progeny of these individuals were grown in grown on an aeroponic chamber and inoculated with *S. medicae* ABS7 rhizobia. The *sobir1-1* homozygotes and the segregating *sobir1-2* homozygotes formed no nodules. Locations of the insertions relative to the gene are displayed in Figure 1.

**Table 1.**
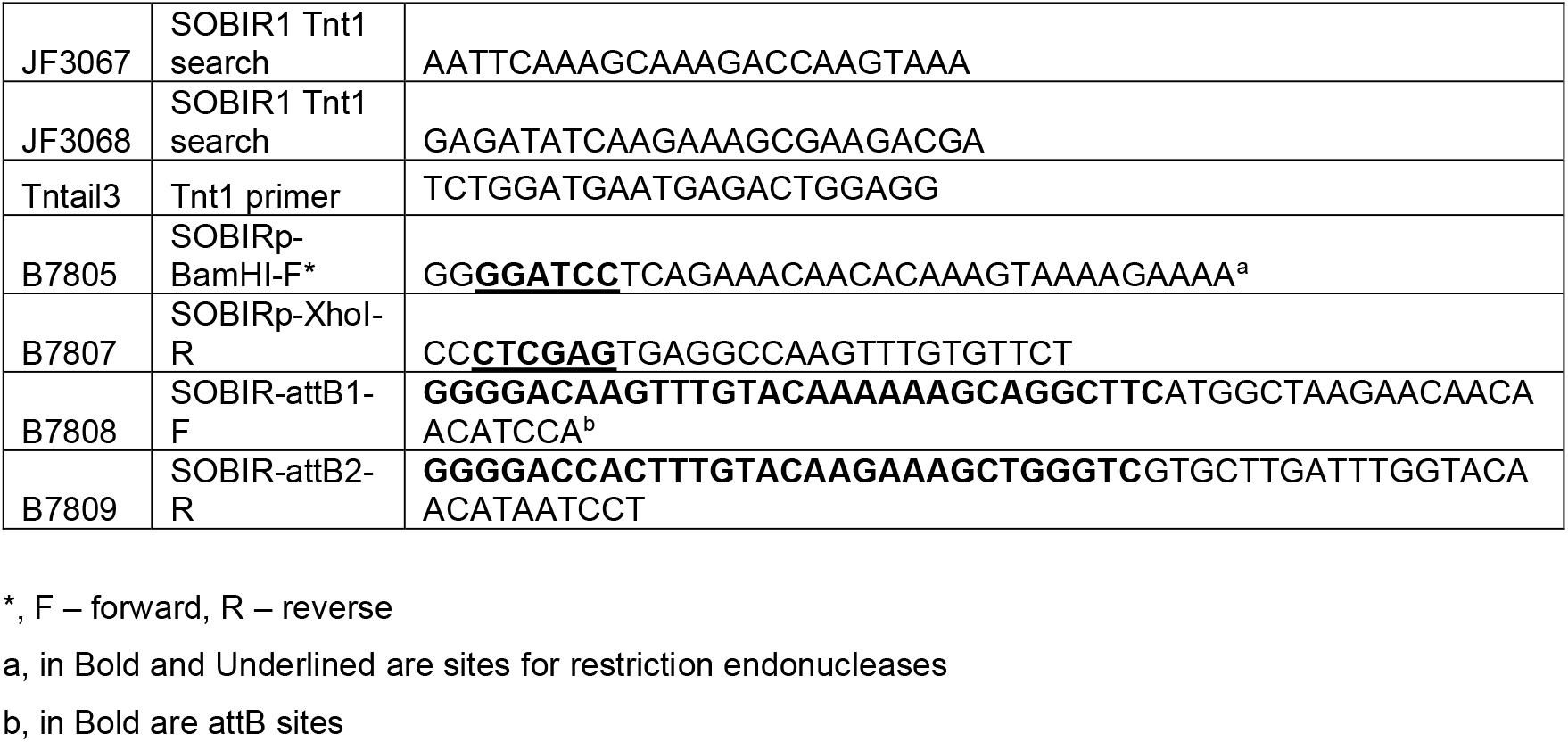
Primers used in this work

### Plant growth conditions and imaging Figure 3B, C, Figure 4 and 5

Wild type R108, *sobir1-1* and *sobir1-2 Medicago truncatula* seeds were scarified with concentrated sulfuric acid for 8 minutes, rinsed multiple times, sterilized with 30% bleach containing a few drops of Tween 20, and rinsed 5 times with sterile water. The seeds were allowed to imbibe in sterile water for 5-7 hours at room temperature, with gentle shaking and multiple changes of water and were placed in a cold room overnight. The next day, the imbibed seeds were plated on 25 mm-deep sterile Petri dishes and placed upside down in the dark at room temperature overnight, to facilitate germination. Germinated seedlings were sown in potting substrate consisting of sterilized turface and vermiculite (2:1; w:w) in Ray Leach Cone-tainers (Stuewe & Sons, Tangent, OR) and were grown in a walk-in growth chamber under a 16-h/8-h light/dark regime, 200 µE m-2 sec-1 light irradiance, 24 °C, and 40% relative humidity. Nodulation experiments were carried out with *Sinorhizobium meliloti* Rm41 (Kondorosi et al., 1977), Sm1021 (Meade et al., 1982) and *S. medicae* ABS7 and WSM419, all carrying the *hemA:LacZ* reporter plasmid pXLGD4 (Leong et al., 1985). Each seedling was inoculated after 7 days of nutrient starvation with 50 ml bacterial suspension at OD600 = 0.03 prepared in one-half-strength B&D solution with 2 mM KNO_3_ (Broughton and Dilworth, 1971).

Plants were harvested at 12- and 20-days post-inoculation. Nodulated root systems were rinsed and photographed using the SMZ18 Nikon stereo microscope equipped with a DS-Ri2 camera and the NIS-Elements BR 5.02.00 software Nikon Instruments, Melville, NY). For histochemical staining, root systems were fixed in 2.5% glutaraldehyde in 100 mM piperazine-N, N′-bis (2-ethanesulfonic acid) (PIPES) buffer, pH7.2 and stained in 5 mM potassium ferricyanide, 5 mM potassium ferrocyanide, 0.08 % 5-bromo-4-chloro-3-indolyl-b-D-galactopyranoside (X-Gal), 0.1% Tween 20 in PIPES, pH 7.2 (Boivin et al., 1990) and processed as previously described (Pislariu et al., 2012).To obtain 50 μm thin sections, stained nodules were embedded in 5% low melting point agarose (GoldBio, St. Louis, MO) and sliced with a PELCO easiSlicer (TedPella Inc., Redding, CA). LacZ-stained whole mounts of root segments and nodule sections were visualized using the Nikon Eclipse 90i Nikon compound microscope equipped with a DS-Fi1 camera and photomicrographs were acquired with the NIS-Elements AR 3.30.02 software (Nikon Instruments, Melville, NY). The same samples were counterstained with the fluorescent DNA dye Syto13 to visualize rhizobia and calcofluor white to visualize cellulose and other beta-glucans in plant cell walls (Life Technologies, Carlsbad, CA) as described by (Haynes et al., 2004). Fluorescently stained nodule sections were observed with an A1 Nikon laser point scanning confocal microscope on an Eclipse Ti inverted microscope equipped with a DS-Fi1 camera. Images were captured using the NIS-Elements AR 5.30.05 software. The Syto13 signal was observed using the GFP laser with excitation at 488 nm and emission at 527 nm. The calcofluor signal was observed using the UV laser with excitation at 405 nm and emission at 440 nm.

### Plant growth and inoculation conditions for AM experiments

For Figure 6, 4-day-old *Medicago truncatula* seedlings were planted in cones filled with a mixture of play sand, filter sand and gravel (ratio 2:2:1), and inoculated with 250 *Glomus versiforme* spores (Müller et al., 2019). Plants were watered twice per week with ½ Hoagland solution containing 20 µM potassium phosphate. Roots were harvested after 4 weeks and stained with WGA-Alexafluor488 to visualize the fungus (Javot et al., 2007). Root length colonization was quantified microscopically using the grid-line method (McGonigle et al., 1990) (n=4-7 plants). Arbuscule numbers were counted in 2mm root segments with a fungal appressorium in the center (n=5 randomly selected segments per plant, three plants per line). Images were taken with a Leica M205 stereomicroscope.

For Figure 7, *M. truncatula* plants were grown as described previously (Ivanov and Harrison, 2024). Briefly, seedlings were planted into 20.5 cm plastic cones filled with a sterile mixture of sand/gravel containing with 100 sterile *Rhizophagus irregularis* spores placed 4 cm below the substrate surface. Plants were grown in a growth chamber under a 16-hour light/22°C and 8-hour dark/20°C regime with 40% humidity. Half-strength Hoagland’s solution containing half-strength nitrogen and 50 μM potassium phosphate was supplied twice a week.

### Plasmid construction

*M. truncatula* accession Jemalong A17 genomic DNA or cDNA was used as a template for PCR amplification of promoter regions and coding sequences of *SOBIR1*. The promoter region (1450 bp) was amplified using oligonucleotides B7805 and B7807 containing sites for restriction endonucleases *Bam*HI and *XhoI*. The amplified DNA fragment was cloned into pENTR L4-R1 containing multiple cloning site using *Bam*HI and *XhoI* (Promega) and T4 DNA ligase (Promega) to obtain pENTR L4-R1 *SOBIR1pro*. The coding sequence without a stop codon was amplified using oligonucleotides B7808 and B7809 containing *att*B recombination sites. The amplified DNA fragment was recombined with pDONR221 using Gateway™ BP Clonase™ II Enzyme mix (ThermoFisher Scientific) to obtain pENTR L1-L2 *SOBIR*. To create an expression vector of translational fusion of *SOBIR* and fluorescent protein *cpVenus*, the pENTR L4-R1 *SOBIR1pro* and pENTR L1-L2 *SOBIR1* were used in the recombination reaction with pENTR R2-L3 *cpVenus* (Lindsey, 2020) and destination vector pK7m34GW BCP-R (Ivanov and Harrison, 2024) using Gateway™ LR Clonase™ II Enzyme mix (ThermoFisher Scientific).

### Confocal laser-scanning microscopy for AM fungi

Transgenic roots showing mCherry fluorescence associated with fungal colonization and expression of AM-specific plasma membrane and periarbuscular membrane marker (*BCP1pro:mCherry-BCP*; BCP-R) were excised into short pieces approximately 3-5 mm in length. The root pieces were then cut longitudinally with a double-edged razor blade and placed on a glass slide with a drop of water with the cut surface facing upwards and covered by a cover slip. Roots sections were observed, and fluorescence was imaged using a Leica TCS-SP5 confocal microscope (Leica Microsystems) with a 20× or 63× water-immersion objective. cpVenus was excited with the argon laser (514 nm) and emitted fluorescence was collected from 529 to 549 nm; mCherry was excited with the Diode-Pumped Solid-State laser at 561 nm and emitted fluorescence was collected from 605 to 630 nm. Images were processed using Leica LAS-AF software versions 2.6.0 (Leica Microsystems), Image J (National Institutes of Health), Adobe Photoshop 2024 version 25.1.0 and Adobe Illustrator 2024 version 28 (Adobe Systems Inc.).

## Acknowledgements

This work was supported by NSF 2139351 (previously NSF 1733470) to Harrison, Frugoli and Pislariu. Mueller and Ivanov were supported by NSF 2139351 (previously NSF 1733470) and a DFG postdoctoral fellowship to Mueller at the time of the experiments.

## Author Contributions

All authors contributed figures and assisted in writing the manuscript. E.S. did the experiments and created/contributed to Figures 1, 2 and 3, C.P. and H.S. to Figures 3, 4, and 5, L.M. to Figure 6 and S.I. to Figure 7.

## References

Albert I, Böhm H, Albert M, Feiler CE, Imkampe J, Wallmeroth N, Brancato C, Raaymakers TM, Oome S, Zhang H (2015) An RLP23–SOBIR1–BAK1 complex mediates NLP-triggered immunity. Nature plants 1: 1–9

Boivin C, Camut S, Malpica CA, Truchet G, Rosenberg C (1990) Rhizobium meliloti Genes Encoding Catabolism of Trigonelline Are Induced under Symbiotic Conditions. Plant Cell 2: 1157–1170

Broughton W, Dilworth M (1971) Control of leghaemoglobin synthesis in snake beans. Biochemical Journal 125: 1075–1080

Cai J, Veerappan V, Arildsen K, Sullivan C, Piechowicz M, Frugoli J, Dickstein R (2023) A Modified Aeroponic System for Growing Plants to Study Root Systems. Plant Methods

Catanzariti AM, Do HT, Bru P, de Sain M, Thatcher LF, Rep M, Jones DA (2017) The tomato I gene for Fusarium wilt resistance encodes an atypical leucine‐rich repeat receptor‐like protein whose function is nevertheless dependent on SOBIR 1 and SERK 3/BAK 1. The Plant Journal 89: 1195–1209

Chaulagain D, Frugoli J (2021) The Regulation of Nodule Number in Legumes Is a Balance of Three Signal Transduction Pathways. International Journal of Molecular Sciences

Choi I-S, Wojciechowski MF, Steele KP, Hopkins A, Ruhlman TA, Jansen RK (2022) Plastid phylogenomics uncovers multiple species in Medicago truncatula (Fabaceae) germplasm accessions. Scientific Reports 12: 21172

Díaz V, Villalobos M, Arriaza K, Flores K, Hernández-Saravia LP, Velásquez A (2025) Decoding the Dialog Between Plants and Arbuscular Mycorrhizal Fungi: A Molecular Genetic Perspective. Genes 16: 143

Domazakis E, Wouters D, Visser RG, Kamoun S, Joosten MH, Vleeshouwers VG (2018) The ELR-SOBIR1 complex functions as a two-component receptor-like kinase to mount defense against Phytophthora infestans. Molecular Plant-Microbe Interactions 31: 795–802

Genre A, Russo G (2016) Does a common pathway transduce symbiotic signals in plant– microbe interactions? Frontiers in plant science 7: 96

Grundy EB, Gresshoff PM, Su H, Ferguson BJ (2023) Legumes Regulate Symbiosis with Rhizobia via Their Innate Immune System. International Journal of Molecular Sciences 24: 2800

Gust AA, Felix G (2014) Receptor like proteins associate with SOBIR1-type of adaptors to form bimolecular receptor kinases. Current opinion in plant biology 21: 104–111

Haynes JG, Czymmek KJ, Carlson CA, Veereshlingam H, Dickstein R, Sherrier DJ (2004) Rapid analysis of legume root nodule development using confocal microscopy. New Phytologist 163: 661–668

Herridge DF, Peoples MB, Boddey RM (2008) Global Inputs of Biological Nitrogen Fixation in Agricultural Systems. Plant Soil 311: 1–18

Ivanov S, Harrison MJ (2024) Receptor-associated kinases control the lipid provisioning program in plant–fungal symbiosis. Science 383: 443–448

Kamphuis LG, Williams AH, D’Souza NK, Pfaff T, Ellwood SR, Groves EJ, Singh KB, Oliver RP, Lichtenzveig J (2007) The Medicago truncatula reference accession A17 has an aberrant chromosomal configuration. New Phytologist 174: 299–303

Kondorosi A, Kiss G, Forrai T, Vincze E, Bánfalvi Z (1977) Circular linkage map of Rhizobium meliloti chromosome. Nature 268: 525–527

Leong SA, Williams PH, Ditta GS (1985) Analysis of the 5’ regulatory region of the gene for delta-aminolevulinic acid synthetase of Rhizobium meliloti. Nucleic Acids Res 13: 5965–5976

Lindsey AR (2020) Sensing, signaling, and secretion: a review and analysis of systems for regulating host interaction in Wolbachia. Genes 11: 813

Madeira F, Park YM, Lee J, Buso N, Gur T, Madhusoodanan N, Basutkar P, Tivey AR, Potter SC, Finn RD (2019) The EMBL-EBI search and sequence analysis tools APIs in 2019. Nucleic acids research 47: W636–W641

Meade HM, Long SR, Ruvkun GB, Brown SE, Ausubel FM (1982) Physical and genetic characterization of symbiotic and auxotrophic mutants of Rhizobium meliloti induced by transposon Tn5 mutagenesis. J Bacteriol 149: 114–122

Ngou BPM, Ding P, Jones JDG (2022) Thirty years of resistance: Zig-zag through the plant immune system. Plant Cell 34: 1447–1478

Pislariu CI, D. Murray J, Wen J, Cosson V, Muni RRD, Wang M. A. Benedito V, Andriankaja A, Cheng X, Jerez IT (2012) A Medicago truncatula tobacco retrotransposon insertion mutant collection with defects in nodule development and symbiotic nitrogen fixation. Plant Physiology 159: 1686–1699

Roy S, Liu W, Nandety RS, Crook A, Mysore KS, Pislariu CI, Frugoli J, Dickstein R, Udvardi MK (2020) Celebrating 20 years of genetic discoveries in legume nodulation and symbiotic nitrogen fixation. The Plant Cell 32: 15–41

Sarrette B, Luu TB, Johansson A, Fliegmann J, Pouzet C, Pichereaux C, Remblière C, Sauviac L, Carles N, Amblard E (2025) Medicago truncatula SOBIR1 controls pathogen immunity and specificity in the Rhizobium‐legume symbiosis. Plant, Cell & Environment 48: 32–50

Schnabel E, Mukherjee A, Smith L, Kassaw T, Long S, Frugoli J (2010) The lss supernodulation mutant of Medicago truncatula reduces expression of the SUNN gene. Plant Physiol 154: 1390–1402

Smith S, Read D (2008) Mycorrhizal Symbiosis (ISBN 978-0-12-370526-6) New York. In. Elsevier

Tadege M, Wen JQ, He J, Tu HD, Kwak Y, Eschstruth A, Cayrel A, Endre G, Zhao PX, Chabaud M, Ratet P, Mysore KS (2008) Large-scale insertional mutagenesis using the Tnt1 retrotransposon in the model legume Medicago truncatula. Plant Journal 54: 335–347

Van Der Burgh AM, Postma J, Robatzek S, Joosten MH (2019) Kinase activity of SOBIR1 and BAK1 is required for immune signalling. Molecular plant pathology 20: 410–422

Vasse J, De Billy F, Camut S, Truchet G (1990) Correlation between ultrastructural differentiation of bacteroids and nitrogen fixation in alfalfa nodules. Journal of Bacteriology 172: 4295–4306

Vasse J, de Billy F, Truchet G (1993) Abortion of infection during Rhizobium meliloti alfalfa symbiotic interaction is accompanied by a hypersensitive reaction. The Plant Journal 4: 555–566

Zhang C, He J, Dai H, Wang G, Zhang X, Wang C, Shi J, Chen X, Wang D, Wang E (2021) Discriminating symbiosis and immunity signals by receptor competition in rice. Proceedings of the National Academy of Sciences 118: e2023738118

Zhang C, Hicks GR, Raikhel NV (2014) Plant vacuole morphology and vacuolar trafficking. Frontiers in Plant Science 5: 476

